# Sex Differences in Motor Unit Properties and Force Steadiness: Insights from Strength-Matched Elbow Flexion

**DOI:** 10.64898/2026.08.06.742881

**Authors:** Parisa Alaei, Kaylee A. Larocque, Changki Kim, Jennifer M. Jakobi

**Affiliations:** Healthy Exercise and Aging Laboratory, University of British Columbia Okanagan, Kelowna, BC, Canada; Faculty of Health and Social Development, University of British Columbia Okanagan, Kelowna, BC, Canada; Department of Kinesiology, College of Education, University of Alabama, Tuscaloosa, AL, USA

**Keywords:** Discharge Rate, Biceps Brachii, Force variability, Intramuscular electromyography, Sex-differences

## Abstract

Sex-related differences in force steadiness are often attributed to maximal strength and motor unit (MU) properties, but their independent contributions remain unclear. This study strength-matched females and males to remove the influence of maximal strength and determine whether MU properties are associated with sex-related differences in force steadiness. Twelve young adults (6 females) were matched for elbow flexion strength (females, 188.6±15.6 N; males, 199.7±24.8 N, p=0.4). Both groups performed submaximal isometric elbow flexion contractions at 2.5%, 5%, 10%, 15%, and 25% MVC. The MU recruitment thresholds (RT), discharge rates (MUDR), and coefficient of variation of interspike intervals (CVISI) were measured from intramuscular fine wire electromyography (EMG) electrodes. Force steadiness was quantified as the standard deviation (SD) and coefficient of variation (CV) of force. Across forces, SD and CV of force did not differ between females and males (p>0.05). Females had a higher recruitment threshold than males (p<0.05). Females had higher MUDR at 15% and 25% MVC (p<0.02), while males were higher at 5% MVC (p=0.02). The CVISI was greater in females (p<0.001) and positively correlated with SD of force (r=0.2) and negatively with CV of force (r=-0.2) in females and males. When strength was matched, sex-related differences in force steadiness were not evident. However, females exhibited higher MU recruitment thresholds, MUDR and CVISI. Despite greater CVISI in females, these differences did not translate into greater force fluctuations, suggesting that individual MU discharge variability is not a primary predictor of force steadiness when maximal strength is controlled.

**NEW & NOTEWORTHY:**

- Strength matching eliminated sex-related differences in elbow flexor force steadiness.
- Females achieved similar force steadiness using higher MU recruitment thresholds and discharge rates, particularly in the short head of the biceps brachii.
- In females, the greater variability in motor unit discharge was not associated with reduced force steadiness.

## INTRODUCTION

Females are typically less steady than males in producing submaximal contractions in both the upper and lower limbs [1–3]. The mechanisms underlying these sex-related differences are not fully understood; however, irrespective of biological sex, isometric force steadiness is influenced by muscular strength and motor unit (MU) activity, both individually and collectively within the pool [4,5]. Stronger individuals are steadier [3,6–8], and because upper limb strength in young adult females is ∼50–70% that of males [9], it is unsurprising that when maximal strength is not controlled females are consistently less steady than males [2,3,6,10]. Thus, strength differences may potentially mask intrinsic differences in force steadiness and underlying MU behavior between females and males.

Without strength matching females generally exhibit higher MU discharge rates (MUDR) and MU discharge variability [2,11,12], the later a factor related to greater force fluctuations [4,13,14]. When strength is matched; however, as shown in small, low-load muscles such as the first dorsal interosseous, females have higher recruitment threshold, MUDR and CV of force [15]. These findings suggest that sex-related differences in MU discharge behavior can persist even when strength is controlled, providing a physiological basis for expecting reduced steadiness in females under strength-matched conditions. Yet, this possibility must be examined in larger, stronger, and functionally relevant proximal muscle groups involved in gross upper limb movements, where neuromechanical demands differ substantially from those of small intrinsic hand muscles.

The elbow flexors, such as the biceps brachii, are larger proximal muscles with greater MU numbers and broader force-generating capacity than small distal muscles [16,17]. These muscles also exhibit distinct recruitment and rate coding strategies compared with the first dorsal interosseous (FDI). Accordingly, it is unclear whether the findings in small distal muscles generalize to larger proximal muscle groups. The biceps brachii comprises the short head (SH), which inserts on the coracoid process and is mechanically more efficient at producing torque at mid-range elbow flexion angles, whereas the long head (LH) crosses the shoulder and contributes across broader joint positions [18]. These architectural and functional differences between muscle heads may further affect MU properties and should be considered when examining force steadiness under strength-matched conditions in females and males.

Despite the historically small number of studies that have directly compared females with males [19,20], recent interest has risen and substantive progress made in characterizing MU properties. Across upper- [15,21] and lower-limb muscles and a range of contraction intensities [22–26], females often exhibit higher MUDR than males. Females also show greater inter-spike interval variability in low-threshold MUs of the biceps brachii [27]. Yet, findings are inconsistent across muscles and contraction intensities, and not observed uniformly at similar relative forces or in comparable muscle groups [12]. The limited and heterogeneous nature of the literature constrains understanding of the factors that contribute to reduced force control in females, highlighting the need for more systematic investigations. Consequently, strength-matching provides a critical method to isolate contributions of the MU to force control in females and males.

Given the importance of strength, the purpose of this study was to determine whether sex-related differences in elbow flexion force steadiness and MU properties are apparent when maximal strength is matched. This approach isolates the contribution of MU properties to steadiness. It was hypothesized that females would remain less steady and exhibit higher MUDR and variability even when strength is matched, reflecting differences in MU properties independent of absolute strength.

## METHODS

### Participants

Twelve young adults (6 females) with no known neurological or musculoskeletal impairments were successfully strength-matched between sexes. Participants who were trained in fine motor control (such as musicians), or who were unable to have their maximal voluntary contraction (MVC) matched to participants of the opposite sex (n=2) were not included in the analysis. All participants provided informed written consent prior to beginning the study, which was approved through the institutional review board (H11-01931) at the University of British Columbia and was conducted in accordance with the Declaration of Helsinki, albeit the study was not registered.

### Experimental Set-up and Procedure

Participants were seated in a custom-designed force dynamometer chair with the right arm abducted at 20°, the shoulder at 0° flexion, and the elbow flexed to 90°. The elbow was supported on a padded rest, and the right hand grasped a manipulandum in a neutral position to record elbow flexion force using a linearly calibrated force transducer (MLP-150, 68kg, sensitivity 32.5 Nm, Transducer Techniques, Temecula, CA, USA). Chest and waist constraints were used to ensure consistent participant positioning and isolation of elbow flexion. Force signals were amplified (×100) (Coulbourn Electronics, Allentown, PA), sampled at 1000 Hz and converted from analog to digital using a Power 1401 (Cambridge Electronic Design, Cambridge, England). Signals were displayed in real time on a 52 cm flat-screen monitor (1920 × 1200 resolution) positioned 1 m in front of the participant at eye level. Data was stored for offline analysis using Spike 2 Version 7.12 (Cambridge Electronic Design, Cambridge, England).

### MVC and Twitch Interpolation

To measure voluntary activation (VA), electrical stimulation (100 µsec pulse width, DS7AH, Digitimer Ltd., Welwyn Garden City, UK) was delivered at 1 Hz percutaneously through two 41cm × 41cm carbon-carbon stimulation electrodes, coated in electrode gel (Parker Spectra 360 Electrode Gel, Parker Lab Inc., Fairfield, NJ, USA) and secured over the proximal and distal aspects of the biceps brachii. The stimulation intensity was increased progressively until a plateau in twitch force amplitude occurred, and then a 10% increase in stimulator output was made to achieve supramaximal intensity. To measure MVC, participants performed three, maximal isometric elbow flexion attempts in the neutral forearm position. Participants were verbally encouraged to pull-up as hard and as fast as possible against the manipulandum. Twitch stimuli were delivered immediately preceding the MVC (resting twitch), during the MVC (interpolated twitch), and immediately following a return to baseline force level (post MVC twitch). Each contraction was separated by ∼2 minutes of rest. The highest MVC value was used to establish an isometric target force for the submaximal isometric tracking tasks.

### Intramuscular Fine-Wires and Force Tracking Tasks

Single MU action potentials were sampled using custom-made bipolar fine-wire electromyography (EMG) electrodes inserted ∼1 cm into the LH and SH (102-µm diameter, California Fine Wire, Grover Beach, CA). Before insertion of the wires, the skin area was exfoliated and cleansed thoroughly with course cleansing pads and 70% ethanol. The fine wire electrodes were inserted intramuscularly with a 27.5 gauge hypodermic needle that was removed once the wire electrodes were in place. The depth and orientation of the wires were altered by gently tugging to draw the wires closer to the surface in an attempt to optimize individual MU recordings. When recordings were less than ideal another electrode was inserted. The reference electrodes were placed on bony prominences of the acromial end of the clavicle and lateral epicondyle of the humerus for the SH and LH, respectively. Signals from the intramuscular fine wire electrodes (California Fine Wire, Grover beach, CA, USA) were high-pass filtered at 10 Hz (NL144, Digitimer Ltd, Welwyn Garden City, UK), pre-amplified at the site of recording with a custom-built amplifier (Don Clarke, University of Windsor, Windsor, ON), hardware amplified (1000 x; NL820A, Digitimer Ltd, Welwyn City, UK), and sampled at 12 kHz (Power 1401 plus, Cambridge Electronics Design, Cambridge, UK).

Force-tracking tasks were done at 2.5%, 5%, 10%, 15% and 25% MVC. The submaximal tracking task involved producing an isometric ramp-up elbow flexion contraction to the targeted force level at 10% MVC/s and subsequently holding the plateau phase for 5 s which was followed by the de-ramp at the same rate. Each trial was separated by 30-60 s of rest to prevent fatigue. Three force tracking tasks were completed for each force level, totaling 15 contractions per participant. The order of contraction intensity was randomized within and between participants. To assess whether a reduction in maximal force-generating capacity occurred as a result of the submaximal tracking protocol the MVC was repeated following the submaximal contractions.

### Data Analysis

Data analyses were performed using Spike2 (version 7.12, Cambridge Electronics Design, Cambridge, UK). Elbow flexion MVC and VA were taken from the highest MVC amplitude prior to the submaximal ramp contractions. VA was calculated using the twitch interpolation technique: VA (%) = (1 - [interpolated twitch/resting twitch]) *100) [28].

Single MU analysis was performed offline (Spike 2 version 7.12; Cambridge Electronics Design, Cambridge, UK). Single MU trains were identified using a template matching algorithm that considers temporal and spatial waveform characteristics of sequential action potentials (Figure 1). Recruitment thresholds (%MVC) were calculated as the level of force at which a MU discharged its first and consistent action potential. Interspike intervals (ISIs), defined as the time between consecutive spikes, were measured during the plateau phase of the contraction. The mean discharge rate for each MU was determined as the reciprocal of the ISIs, averaged across the plateau phase. Additionally, the coefficient of variation of the ISIs (CV ISI) was calculated as the standard deviation divided by the mean ISI to quantify MU discharge variability. Force steadiness was evaluated offline for the plateau phase of each contraction, quantified as both the standard deviation (SD) and the coefficient of variation (CV) of force, with CV calculated as SD divided by the mean force.

**Figure 1.**
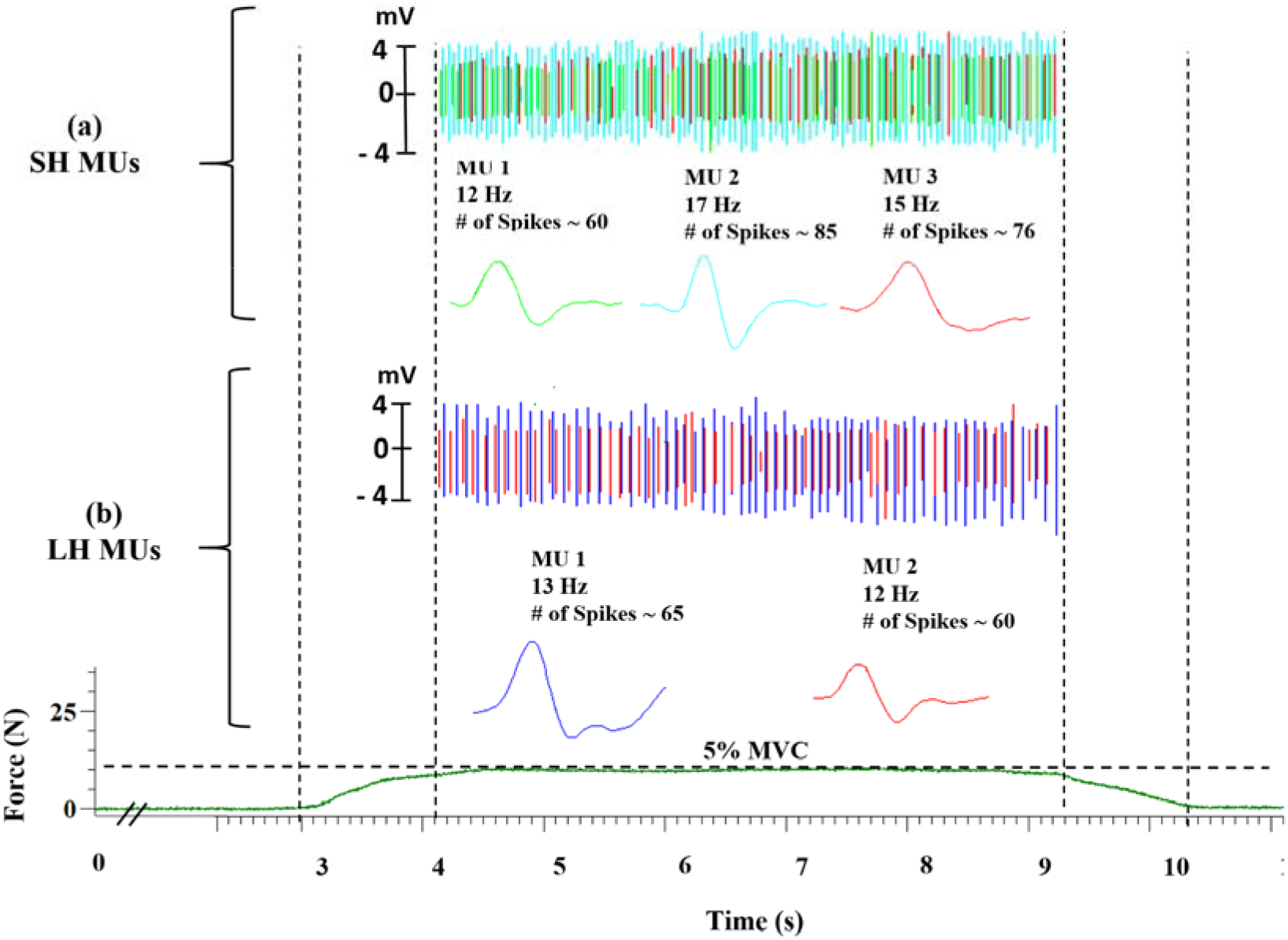
Representative data from 5% MVC of a female. MU action potential trains for the (a) short (SH) and (b) long (LH) head of the biceps brachii as well as individual action potentials from the spike trains during the plateau phase of a 5% isometric elbow flexion contraction. MU, Motor Unit; SH, Short Head; LH, Long Head; #, number.

### Statistical Analyses

Statistical analyses were performed using IBM SPSS Statistics version 27 (IBM, Armonk, NY, USA). Normality was assessed using the Shapiro–Wilk test. As the data were not normally distributed, a Kruskal–Wallis test was used to examine differences in outcome measures across sex, muscle, and force level. When the Kruskal–Wallis test indicated a significant overall difference, pairwise comparisons were performed using Mann–Whitney U tests. Spearman’s correlation analyses were conducted separately for males and females to examine relationships between SD and CV of force and CVISI. Statistical significance was set at P ≤ 0.05, and data are reported as mean ± SD.

## RESULTS

Six young females (22.0 ± 2.5 years; 68.7 ± 6.4 kg; 167.7 ± 5.6 cm) and six young males (24.0 ± 3.0 years; 60.9 ± 9.6 kg; 168.7 ± 10.8 cm) were successfully matched for strength (females, 188.6 ± 15.6 N; males, 199.7 ± 24.8 N; t (10) = 0.9, p = 0.4) and VA scores did not differ between the sexes (females, 99.2 ± 1.5%; males, 97.7 ± 2.02 %; t (9) = 1.44, p = 0.2). Motor unit properties A total of 411 MUs were recorded, with no differences in the number of potentials between the LH and SH or between females and males (p>0.5; Table 1). An average of 3.3 ± 1.8 MUs were recorded per trial, and did not differ between contraction levels (P > 0.05) with recordings in females 3.2 ± 1.9 and males 3.3 ± 1.7 also not differing t (141) = −0.299, p = 0.8.

**Table 1.** Number of recorded MUs by muscle and force level in males and females.

| *Force Level* | *Females* |  |  | *Males* |  |  |
| --- | --- | --- | --- | --- | --- | --- |
|  | *LH* | *SH* | *Total* | *LH* | *SH* | *Total* |
| 2.5% | 18 | 10 | 28 | 15 | 16 | 31 |
| 5% | 20 | 15 | 35 | 23 | 20 | 43 |
| 10% | 22 | 17 | 39 | 26 | 20 | 46 |
| 15% | 35 | 20 | 55 | 31 | 20 | 60 |
| 25% | 21 | 21 | 42 | 18 | 14 | 32 |
| ***Total*** | 116 | 83 | 199 | 113 | 90 | 212 |

LH, Long head of biceps brachii; SH, Short head of biceps brachii

Females had higher recruitment threshold than males at all force levels above 2.5% MVC (5%: H _(1)_ = 6.03, p = 0.01; 10%: H _(1)_ = 3.96, p = 0.04; 15%: H _(1)_ = 17.96, p < 0.001; 25%: H _(1)_ = 6.8, p = 0.009) (Figure 2A). The recruitment threshold at 25% MVC was significantly higher than all other force levels (p < 0.05), and 2.5% and 5% MVC was lower than 10%, 15%, and 25% MVC (p < 0.05). Females also exhibited higher recruitment threshold than males in both the LH (H _(1)_ = 8.9, p = 0.003) and SH (H _(1)_ = 19.4, p < 0.001) (Figure 2B).

**Figure 2.**
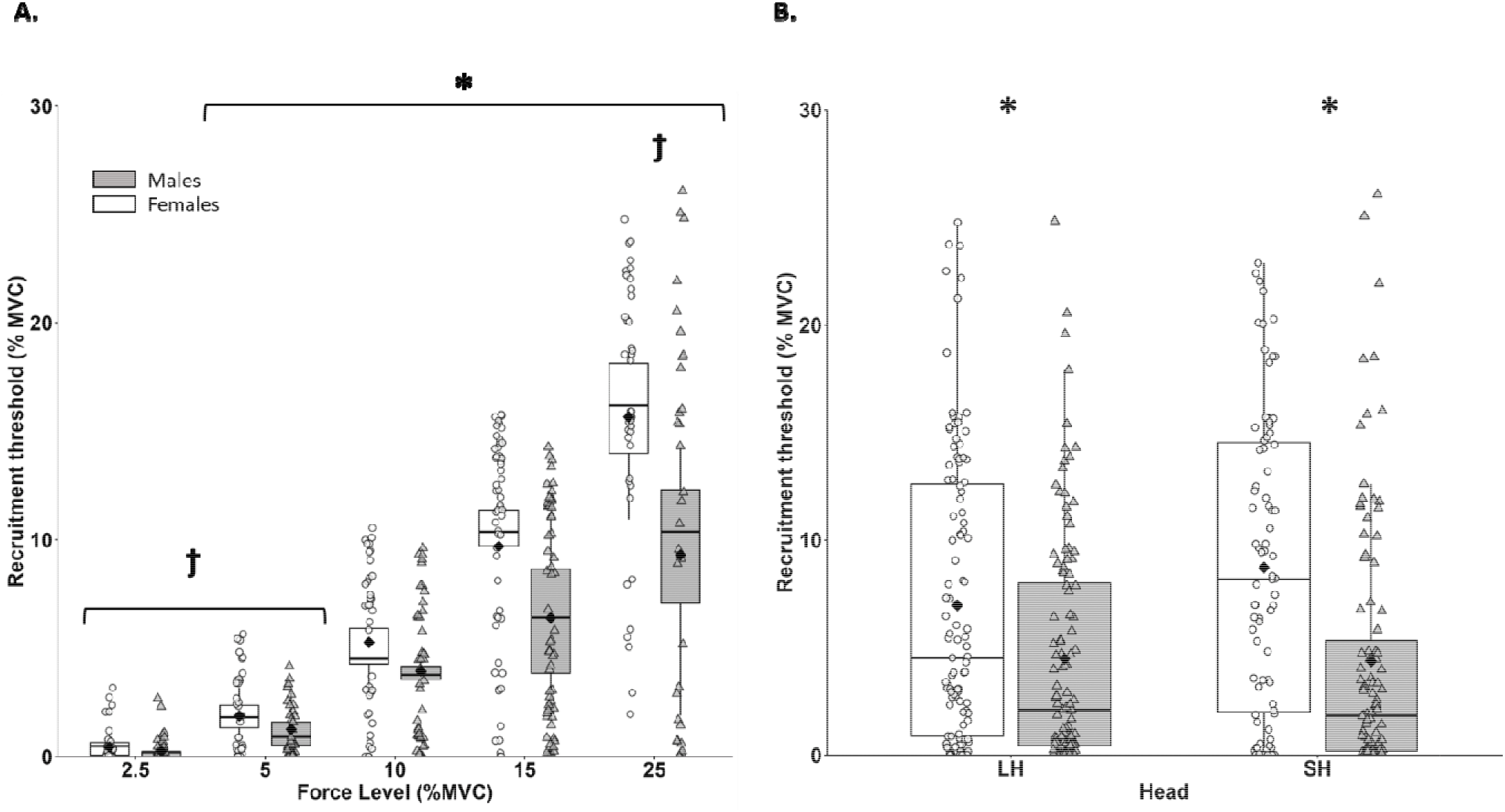
Box plots presenting recruitment threshold of the biceps brachii between females (open box plots; open circles individual data) and males (filled box plots; filled triangles individual data) (A) across the submaximal force levels and (B) between long head and short head. *, females differ from males; IZ, differs from 10%, and 15%. MVC, maximal voluntary contraction; LH, Long Head; SH, Short Head. Black diamonds are mean values. Number of motor units recorded: females = 199; Males = 212

Females had higher MUDR than males at 15% (H _(1)_ = 7.8, p = 0.005) and 25% (H _(1)_ = 5.2, p = 0.022) of MVC. Males were higher at 5% (H _(1)_ = 5.2, p = 0.022) (Figure 3A). MUDR was higher at 15% and 25% than other force levels (p < 0.05) for both males and females. In the LH (H _(1)_ = 1.8, p = 0.2) and SH (H _(1)_ = 3.1, p = 0.08) MUDR was not significantly different between females (LH: 16.12 ± 5.0; SH: 16.01 ± 4.3 %MVC) and males (LH: 14.73 ± 3.2; SH: 14.8 ± 2.6 %MVC).

**Figure 3.**
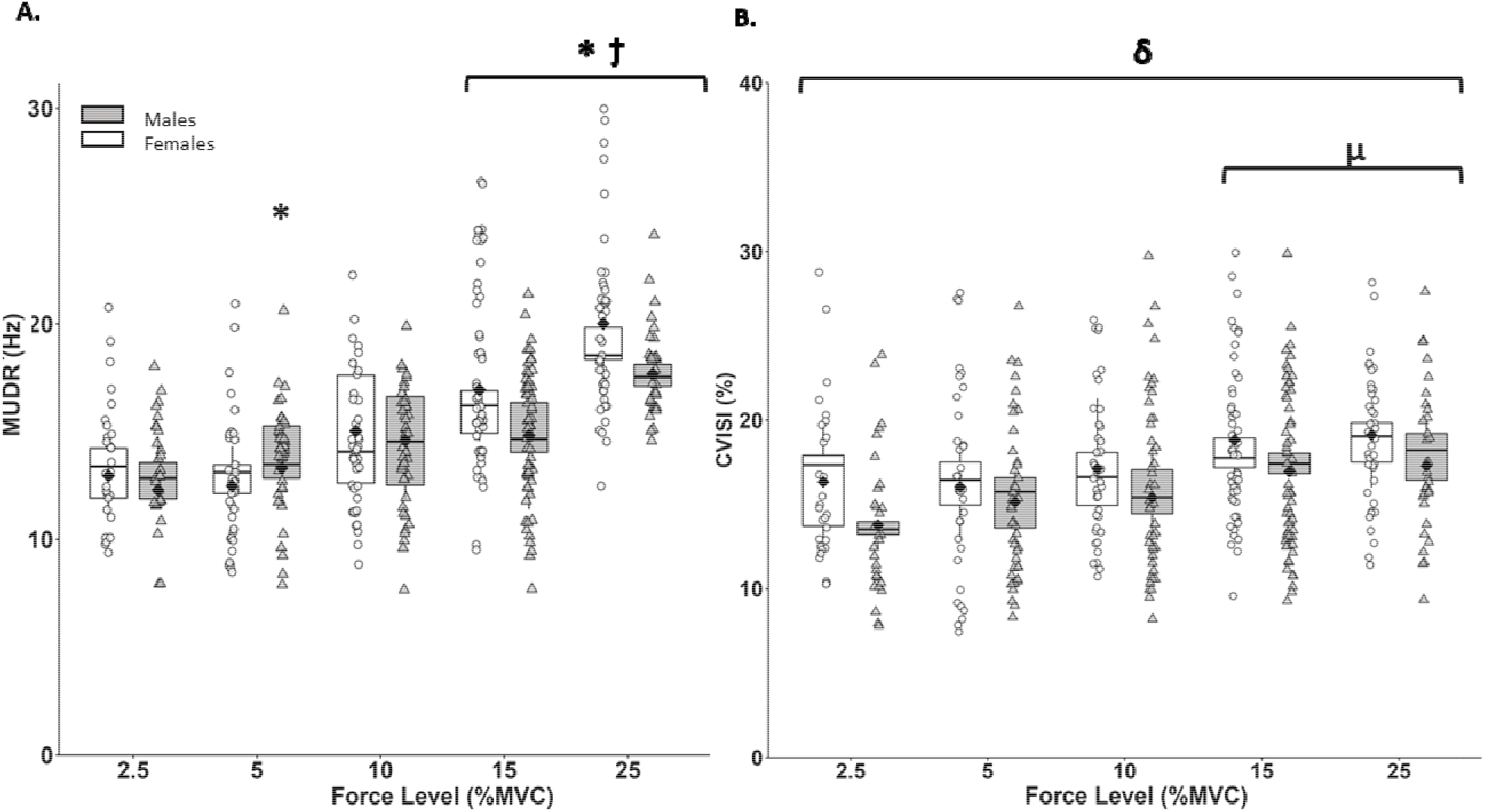
Box plots of (A) MUDR and (B) CVISI for females (open boxplots; open circles individual data) and males (filled boxplots; filled triangles individual data) across all submaximal force levels. *, females differ from males; δ, females are higher than males across all forces; IZ, differs from other force levels; μ, differs from 2.5 and 5% MVC. MVC, maximal voluntary contraction; MUDR, motor unit discharge rate; CVISI, coefficient of variation of the inter-spike intervals. Black diamonds are mean values. Number of motor units recorded: females = 199; Males = 212

A significant main effect of sex was observed, with females exhibiting higher CVISI than males (H _(1)_ = 12.665, p < 0.001) across all force levels and muscles. At 2.5% force, CVISI was higher in females (H _(1)_ = 3.7, p = 0.05), while no sex-related differences were observed at 5% (H _(1)_ = 1.34, p = 0.3), 10% (H _(1)_ = 1.6, p = 0.2), 15% (H _(1)_ = 3.5, p = 0.06), or 25% (H _(1)_ = 1.8, p = 0.2) force levels (Figure 3B). In the SH, females (18.4 ± 4.3%) had higher CVISI than males (15.8 ± 4.9%) (H _(1)_ = 15.2, p < 0.001) but in the LH, there was no significant difference between females (17.1 ± 4.8 %MVC) and males (16.2 ± 4.7%) (H _(1)_ = 1.6, p = 0.2). CV ISI was higher at 15% and 25% than 2.5% and 5% force levels (p < 0.05).

### Force Steadiness

The SD of force did not differ between females and males across all submaximal force levels: 2.5% (H _(1)_ = 1.05, p = 0.3), 5% (H _(1)_ = 0.19, p = 0.7), 10% (H _(1)_ = 0.73, p = 0.4), 15% (H _(1)_ = 0.4, p = 0.5) and 25% (H _(1)_ = 1.43, p = 0.2) of MVC. Between forces levels, the SD of force at 2.5% MVC was lower than at 10%, 15%, and 25% MVC (p < 0.05), and 5% MVC was lower than 15% and 25% MVC (p < 0.05), and 25% MVC was significantly higher than all other force level (p < 0.05) (Figure 4A). Similarly, the CV of force did not differ between females and males at 2.5% (H _(1)_ = 0.04, p = 0.8), 5% (H _(1)_ = 0.96, p = 0.3), 10% (H _(1)_ = 2.7, p = 0.1), 15% (H _(1)_ = 1.4, p = 0.2) and 25% (H _(1)_ = 3.2, p = 0.1) of MVC. Irrespective of sex, CV of force was significantly higher at 2.5% and 5% MVC than 10%, 15%, and 25% (p < 0.05) (Figure 4B).

**Figure 4.**
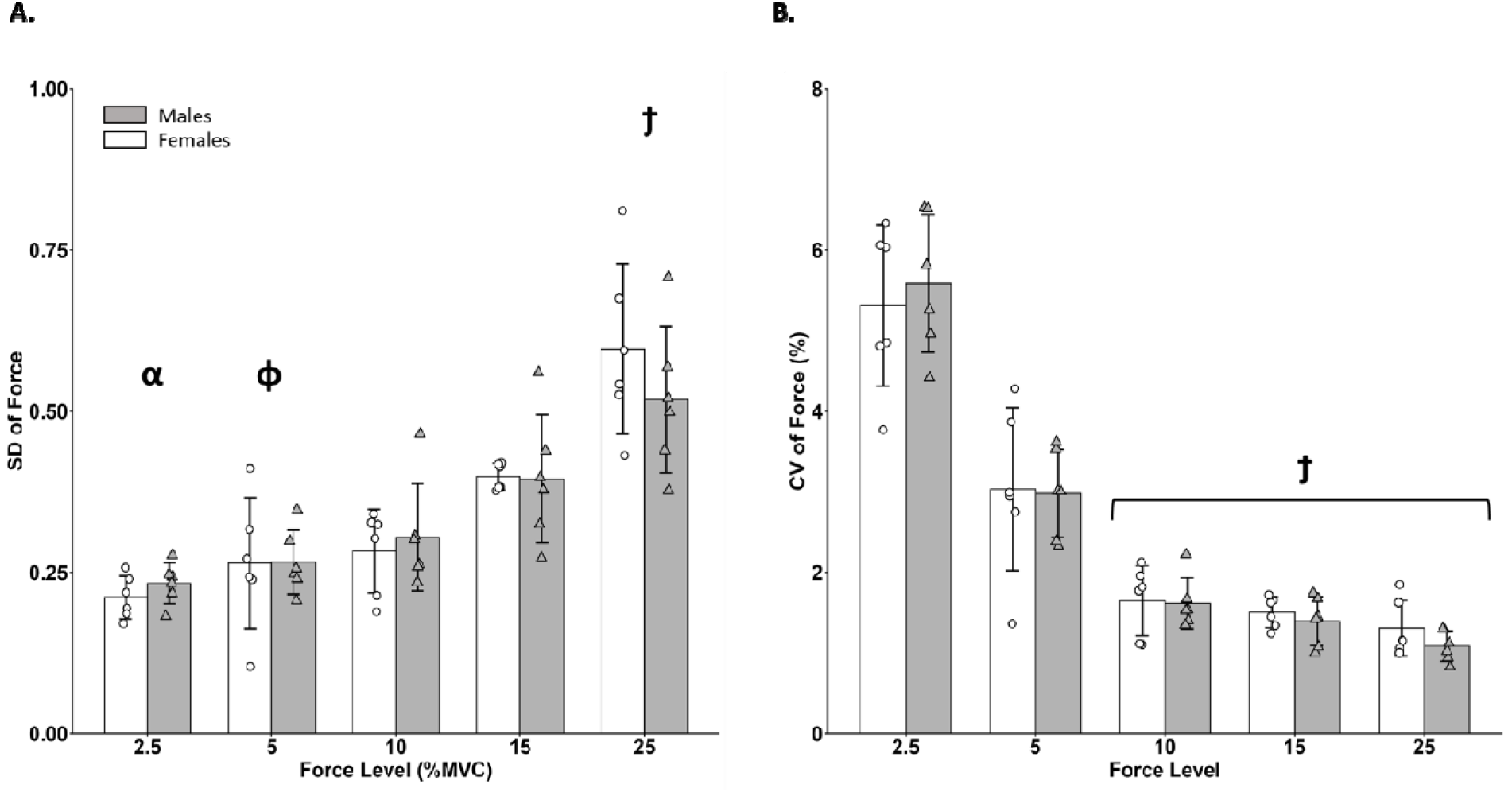
(A) SD of force and (B) CV of force between females (open bars; open circles individual data) and males (filled bars; filled triangles individual data) across all submaximal force levels. IZ, differs from other force levels; α, differs from 10%,15%, and 25%; IZ, differs from 15% and 25%. MVC, maximal voluntary contraction; SD, Standard Deviation; CV, coefficient of variation. Values are mean ± SD. Number of participants: females = 6; Males = 6

### Correlation analysis

Spearman correlation analysis demonstrated a significant positive correlation between CV ISI and SD of force in both females (r = 0.18, p = 0.009) and males (r = 0.23, p = 0.001) (Figure 5a), and there was also a significant negative correlation between CV ISI and CV of force in both females (r = - 0.19, p = 0.006) and males (r = - 0.25, p < 0.001; Figure 5b).

**Figure 5.**
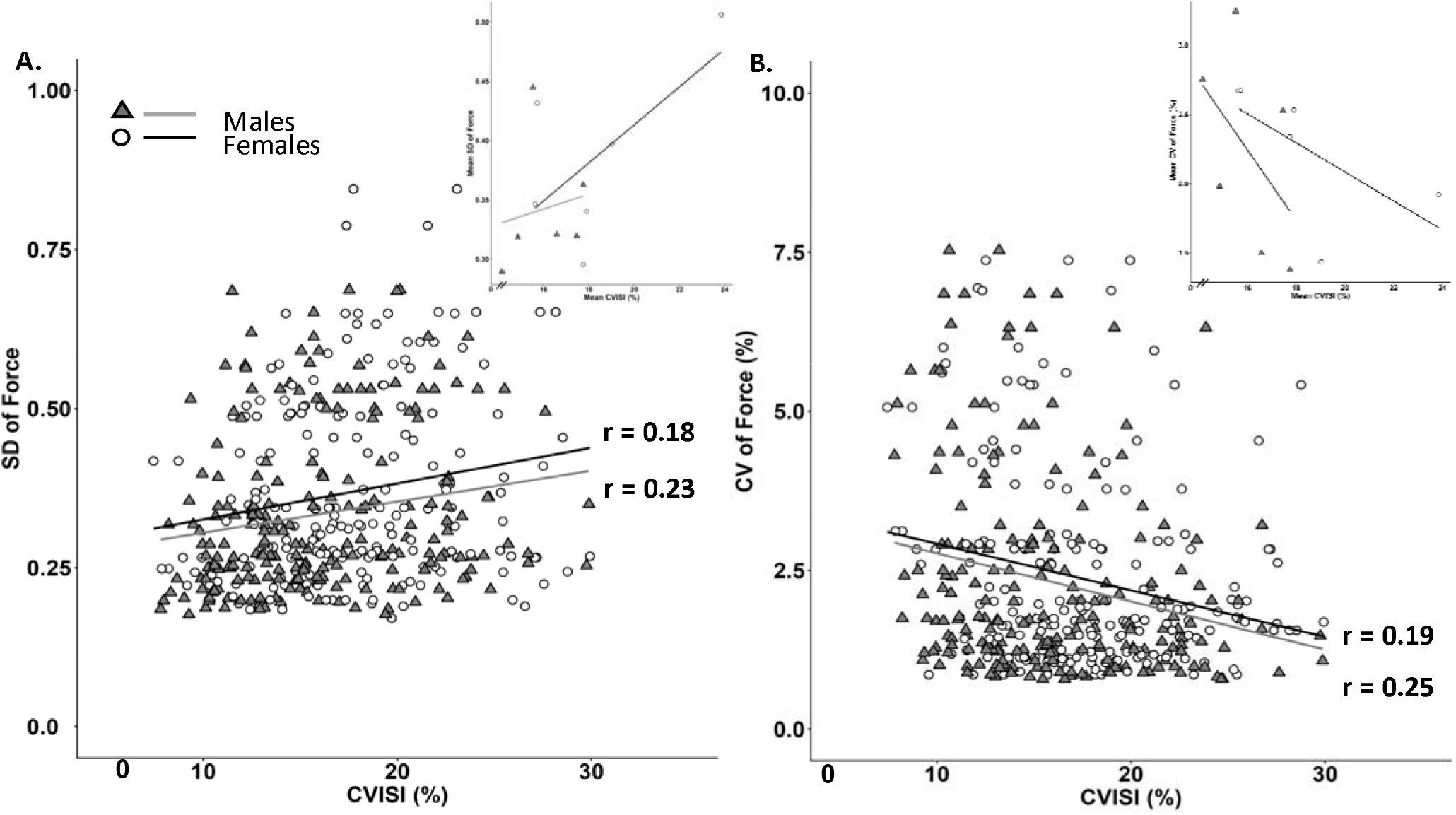
Correlation between (A) CVISI and SD of force and (B) CVISI and CV of force across all force levels in both females (open circles) and males (filled triangles). The inset plot is the average value of CVISI and CV of force per participant at each force level. CVISI, coefficient of variation of interspike interval; SD, Standard Deviation; CV, coefficient of variation. The inset plots in each panel show the same correlations using participant mean values averaged across force levels. Number of motor units recorded: females = 199; Males = 212

## DISCUSSION

The objective of this study was to mitigate the influence of absolute strength when examining sex-specific differences in elbow flexor force steadiness and MU properties in young females and males. The novelty and strength of this study is matching the isometric elbow flexion MVC, between young females and males. When elbow flexion strength is matched force steadiness did not differ between sexes across low force levels of 2.5-25% MVC. Females exhibited higher recruitment thresholds and MUDRs than males. Additionally, the CVISI was also higher in females, particularly in the SH, which is the primary head contributing to force production in a neutral forearm position. Correlational analysis showed that CVISI was positively associated with SD of force and negatively with CV of force. Taken together, strength-matched females and males show comparable force steadiness at low contraction intensities despite females exhibiting higher MU recruitment thresholds, discharge rates, and discharge variability. MU discharge variability was positively correlated with SD of force and negatively associated with CV of force in both sexes, suggesting independent variability in MUs might not be a primary factor contributing to force steadiness.

Recruitment thresholds were higher in females than males, and MUDR was elevated at 15% and 25% MVC, consistent with prior high density surface EMG (HDsEMG) studies reporting sexl1lspecific differences in MU behavior [15,29–32]. Although these earlier findings were partly attributed to methodological factors of HDsEMG including the effects of subcutaneous tissue, impedance, and decomposition algorithms that may bias estimates in females (Taylor et al., 2022), these factors are minimized by the use of indwelling electrodes. This suggests that higher recruitment thresholds observed in females are not simply an artifact of HDsEMG but likely reflect intrinsic MU properties.

Physiologically, it is likely that there is a greater proportion and area of type I MUs in females [33,34]. Type I MUs generate less force, so females must recruit additional and relatively higherl1lthreshold units to achieve equivalent force output, which elevates average recruitment thresholds. At recruitment, these MUs produce less force which necessitates higher firing rates, contributing to increased MUDR [35,36]. These strengthl1lmatched findings align with previously reported higher MUDRs across contraction intensities [2,12,21–24] and may also reflect previously proposed contributions from greater persistent inward currents in females, which enhance motoneuron excitability [30].

Methodological factors may still play a minor role as females typically exhibit greater subcutaneous fat thickness, and although this was not measured, similar needle insertion depth could have shifted sampling toward more superficial muscle regions in females. Although this remains speculative, deeper regions of some grossl1lmovement muscles contain relatively more type I fibers [37], and thus the sample of MUs in females may have been from regions more superficial and more likely to yield higherl1lthreshold (Type II) MUs. Because depthl1ldependent fiberl1ltype distribution in the human biceps brachii is not well established, such methodological interpretations should be viewed cautiously, and future studies using ultrasoundl1lguided electrode placement or insertionl1ldepth normalization may help clarify these possibilities.

Females in this study also exhibited higher CVISI than males, especially in the SH. This pattern aligns with findings from Lecce et al (2024), who similarly reported greater discharge variability in the female biceps brachii. One possible explanation also involves the intrinsic neural factors as the smaller motor neurons in females have higher input resistance [38,39], meaning equivalent synaptic currents can produce larger fluctuations in membrane potential, increasing susceptibility to synaptic noise and elevating discharge variability. Although plausible, this interpretation remains speculative, and alternative mechanisms, including potential hormonal influences, cannot be excluded. Hormonal fluctuations have been proposed as a potential contributor to sex-related differences in MU discharge behaviour [11,40], potentially through effects on motoneuron excitability, synaptic transmission, or central drive; however, ovarian hormone-specific influence cannot be directly assessed in the present study as circulating hormone concentrations were not measured and ovulation was not confirmed.

The difference in MU discharge variability between the SH and LH may reflect anatomical distinctions in elbow flexion mechanics. The SH inserts on the coracoid process of the scapula, providing a mechanical advantage that enables more efficient torque production at 90° of elbow flexion [18]. In this study contractions were performed at 90°, and the SH may have operated closer to its optimal length-tension relationship, reducing drive requirement and leading to later recruitment relative to the LH. Under these mechanically advantageous conditions, the influence of synaptic noise on motor neuron output may become more pronounced. In females, greater fluctuations in synaptic input to the motor neuron pool, combined with the reduced activation requirements of the SH, may increase susceptibility to MU variability, contributing to the higher CVISI observed in the SH. The link between muscle head-specific mechanics and sex-related differences in synaptic input remain to be established, yet these findings reflect compartment-specific neuromechanical organization within the biceps brachii.

Sex-related differences in force steadiness have been attributed to males’ greater absolute strength [1,12,41], and this study intentionally matched isometric elbow flexion force between sexes to minimize strength as a confounding factor. Contrary to our hypothesis, no sex-related differences in force steadiness were observed under strength-matched conditions; both SD and CV of force did not differ between females and males at low to moderate forces. These findings suggest that increased MU discharge variability observed in females does not necessarily translate to reduced elbow flexor force steadiness when the combined action of the biceps brachii, brachialis, and brachioradialis are contributing to force output. Instead, the absence of sex-specific differences in steadiness under strength-matched conditions reinforces the growing body of literature showing that absolute strength is a critical determinant of force steadiness, and that previously reported sex-related differences likely reflect males’ greater maximal strength rather than inherent sex-specific impairments in female force control. Notably, although force steadiness was equivalent between sexes, differences in MU behavior persisted, suggesting that females and males may rely on distinct neuromuscular strategies to achieve similar levels of force steadiness.

To further examine the association between MU behaviour and force steadiness, correlation analysis were conducted between MU CVISI and the SD and CV of force. Spearman correlation analysis revealed a significant positive correlation between CVISI and the SD of force, consistent with prior work showing that greater discharge variability contributes to larger absolute force fluctuations [7,13,14,42,43]. In contrast, earlier studies reported a positive correlation between CVISI and CV of force [14,44], whereas a negative correlation was observed in this study. This likely reflects a mathematical consequence of normalization. Conditions with greater MU discharge variability also produced higher mean forces, so although absolute force fluctuations were larger, they represented a smaller proportion of the total force, yielding lower relative force variability. It is also possible that variability at the single MU level does not directly contribute to differences in whole-muscle force steadiness because the pooled activity of many MUs, combined with common synaptic input to the motor neuron pool, may buffer the influence of individual discharge variability.

When strength was controlled, sex-related differences in SD and CV of force were no longer evident. Nonetheless, CVISI remained significantly associated with force steadiness in both females and males. This raises an important question: if females exhibited higher CVISI, why did force steadiness not differ between sexes? One explanation is that the pronounced discharge variability in females was most evident in the SH of the biceps brachii, whereas force steadiness reflects the combined output of the entire elbow-flexor group, potentially attenuating the impact of localized differences in discharge variability. It is also possible that between-participant differences in CVISI do not predict differences in force steadiness. Future studies should therefore examine longer sustained submaximal contractions in order to use cross-correlation analysis to evaluate the within-trial temporal relationship between fluctuations in force and ISI variability, thereby quantifying the temporal lag between these signals and whether it differs by sex. Another factor that cannot be overlooked is the small sample of strength-matched participants. Achieving strength-matched cohorts in young adults is inherently challenging, and the females in this study self-reported consistent exercise training and were stronger than the typical university-aged female, whereas the males were smaller and lighter than the typical male participant commonly recruited in motor control research [21,27,36]. Differences in training status and physical activity could therefore have influenced force control and biased the interpretation of sex-related differences. Future studies should quantify physical activity patterns rigorously and move beyond standardized questionnaires by assessing sport- and training-specific adaptations, which are known to increase MUDRs, enhance rate of torque development, and improve force steadiness [45–47].

Although the participant sample was small, the total number of MUs recorded (∼ 400) is substantial relative to prior indwelling EMG studies [48–51], and sufficient for characterizing MU properties. Importantly, and unlike many HDsEMG studies MU yield was similar between females and males across both heads of the biceps brachii at all force levels, indicating that females did not contribute fewer MUs to the analysis; an occasional concern in HDsEMG [32]. Collectively, these findings demonstrate that despite females exhibiting higher MU discharge variability, force steadiness remains comparable between sexes when elbow flexor strength is matched.

## CONCLUSIONS

When absolute elbow flexor strength was matched between young females and males, sex-related differences in force steadiness were not evident across low to moderate contraction intensities, even though females exhibited greater MU discharge variability, particularly in the SH of biceps brachii. Females also demonstrated higher MU recruitment thresholds and discharge rates, consistent with previous reports of sex-related differences in MU behavior. Importantly, the greater MU discharge variability observed in females did not translate into differences in force steadiness, indicating that variability at the single MU level is not associated with force steadiness when maximal strength is controlled. Rather, the neuromuscular system may offset sex-related differences in individual MU behaviours through muscle synergists or the pool of MUs; however, these have not been systematically assessed. Overall, these findings highlight that sex-related differences in MU properties do not necessarily manifest as differences in force steadiness when maximal strength is matched, underscoring the importance of system-level control in the expression of force variability.

## ACKNOWLEDGEMENTS

We sincerely thank all the participants for their time and dedication, which were vital to the success of this research.

## AUTHOR CONTRIBUTIONS

P.A: Data curation and analysis, statistics, writing original draft, and revising. K.A.L: Data collection, methodology, data curation and analysis. C.K: review & editing document. J.M.J: Supervision, conceptualization, data collection, methodology, statistics, experimental resources, writing & editing document. All authors have read and agreed to the published version of the manuscript.

## DATA AVAILABILITY

The data supporting the conclusions of this article will be made available by the authors, upon reasonable request, without undue reservation.

